# RFEX: Simple Random Forest Model and Sample Explainer for non-Machine Learning experts

**DOI:** 10.1101/819078

**Authors:** D. Petkovic, A. Alavi, D. Cai, J. Yang, S. Barlaskar

## Abstract

Machine Learning (ML) is becoming an increasingly critical technology in many areas. However, its complexity and its frequent non-transparency create significant challenges, especially in the biomedical and health areas. One of the critical components in addressing the above challenges is the *explainability* or *transparency* of ML systems, which refers to the *model* (related to the whole data) and *sample explainability* (related to specific samples). Our research focuses on both model and sample explainability of Random Forest (RF) classifiers. Our <u>RF ex</u>plainer, RFEX, is designed from the ground up with non-ML experts in mind, and with simplicity and familiarity, e.g. providing a one-page tabular output and measures familiar to most users. In this paper we present significant improvement in *RFEX Model explainer* compared to the version published previously, a new *RFEX Sample explainer* that provides explanation of how the RF classifies a particular data sample and is designed to directly relate to RFEX Model explainer, and a RFEX Model and Sample explainer case study from our collaboration with the J. Craig Venter Institute (JCVI). We show that our approach offers a simple yet powerful means of explaining RF classification at the model and sample levels, and in some cases even points to areas of new investigation. RFEX is easy to implement using available RF tools and its tabular format offers easy-to-understand representations for non-experts, enabling them to better leverage the RF technology.

## 1. Introduction

Machine Learning (ML) is becoming an increasingly critical technology in many areas (biomedicine, health, autonomous driving cars, business, loan approvals, law enforcement, distribution of government and health services, news filtering etc.). However, its complexity and its frequent “non-transparency” create challenges in ensuring its proper, ethical, unbiased, technically and legally transparent (explainable) operations. This is especially of importance in the biomedical and health areas where ML may have direct and serious impact on human life and related products and services as well as on general adoption of these technologies [1,2,3]. Issues related to the use and problems with ML (including trust of users and adopters) are gaining not only attention from technical and academic community but also from business, public media, as well as regulatory and political organizations. One of the critical components in addressing the above challenges is the *explainability* or *transparency* of ML systems. At a high level, explainability properties of ML systems allow humans (experts and non-experts) to gain insights into how ML systems make their decisions. ML explainability can be *model* based, where it offers insights on how the ML system works as a whole on a collection of data (e.g. whole training database) and *sample* based, where it offers insights into how the ML system classifies a specific data sample. The latter has been shown to be critical in determining user trust of non-ML experts who are often the key adopters of this technology [10]. ML explainability is also an essential component of powerful “user in the loop” model of understanding, tuning, debugging, deploying and maintaining ML systems [4,26]. ML explainers can be *agnostic* to ML methods or ML specifics. ML systems that are explainable may achieve many benefits such as: increasing user trust; improvement in quality control and maintenance; legal transparency, and they may even offer new insights into the analyzed domain. The need for explainability does not exclude usefulness of black box models since they are always tried first and serve among other things to point to ultimate achievable accuracy and as such are part of the adoption decision process [9]. Note that some ML methods like Neural Networks and Deep Learning are inherently difficult to explain, while some like tree-based ones are more amenable to explanations. The academic community is addressing this issue with more research, dedicated programs and collaborations. A number of dedicated conferences and workshops on this topic have been held e.g. [8] including at PSB [6,7]. Government funding agencies are promoting research in this area (e.g. DARPA XAI initiative [4]) and think tanks and even companies are producing guidance for better and more ethical use of ML and Artificial Intelligence (AI), notable being Asilomar AI Principles [5] whose recommendations have recently been endorsed by CA Legislature. One of the most well-known ML explainer is LIME [11,12] which is agnostic to ML algorithms and provides explanation information based on analysis and approximations using “black box” ML system response to a set of individual samples. ML (and its explainability) has also been used in genetic research to automatically develop information for classification for ontology databases and for quality control of data extracted by gene sequencing [21,22, 27]. “User in the loop” paradigm for improvement of ML systems are gaining attention [4] with a good example in [26] where authors demonstrate benefits of this approach (they call it “explanatory debugging”) to improving the quality of ML solution and increasing user trust, with explainability as its very important component. Jointly with Stanford Bioengineering we have been working for a number of years in applying ML to biomedical problems [16,17,18] and naturally we got involved in ML explainability, especially of well known Random Forest (RF) ML method [13]. We originally developed <u>RF Ex</u>plainability Enhancement pipeline (RFEX) as RF Model explainer and applied it to Stanford FEATURE data [16] where we also performed usability experiment with 13 users and demonstrated that RFEX increased their understanding of RF classification. In this paper we present several key contributions: a) significant improvements in *RFEX Model explainer* compared to the previously published version [16]; b) a new *RFEX Sample explainer* which provides simple explanation of how RF classifies a particular data sample and is designed to directly relate to RFEX Model explainer; and c) an RFEX Model and Sample explainer case study on the data and findings from J. Craig Venter Institute (JCVI) and Allen Institute for Brain Science [21,22,27] and on the Stanford FEATURE data described in [16].

## 2. Our approach to <u>R</u>andom <u>F</u>orest Model and Sample <u>Ex</u>plainability - RFEX

A number of researchers have leveraged RF’s ability to determine feature importance for addressing the explainability challenge or to transform RF into sets of rules, as covered in [16] and also applied in e.g. [21, 22]. Our RFEX approach is novel in the following: a) it goes far beyond simple ranking of features and provides many other measures to enhance RF explainability (e.g. tradeoffs vs. accuracy, ranking of feature combinations, feature interactions via feature cliques); b) provides RF Model as well as Sample explainers; and c) offers a very simple output report with components familiar to our target users, who are often non-ML experts. RFEX is designed to work only with RF, hence it is a direct method. All estimates of accuracy (e.g. in using subsets of features) are direct e.g. computed using RF “engine” hence are not approximated. RF is widely used, powerful (as evaluated in [14]), well supported with tools (e.g. [15]) and inherently explainable. It is based on sets of *ntree* decision trees (forest of trees) voting together to determine sample classification. The parameter *cutoff* is chosen for voting threshold and determines RF sensitivity. RF has a built in approximately 1/3 cross validation error estimate (OOB) in that samples used for training of each tree are not used for accuracy estimation. In each tree, RF makes decisions on one feature at a time (optimal one from randomly selected *mtry* subset of all features, with replacement) effectively producing box like decision surfaces parallel with feature axes. This is amenable to explanations and consistent with applying range tests on individual features, similarly as common practice of evaluating individual tests in biomedicine. For the main classification accuracy measure, to be more precise in case of imbalanced data, we use standard F1 score computed as 2*(recall*precision)/(recall + precision). RF has several built in feature importance (ranking) measures and we use the MDA (Mean Decrease in Accuracy) perturbation measure [13, 15]. We address the most common case of binary classification.

Specific RFEX design goals were developed in consultation and conversations with our target users namely domain experts (but not ML experts) attempting to use ML technologies and were initially evaluated in [16] with 13 users. RFEX design goals include: a) *user centered design:* we specifically focused on developing explainers that are driven by the specific needs of our intended users; and b) *simplicity and familiarity*: we enforced simplicity and familiarity with common ways and measures our users analyze the data such as medical tests e.g. we enforced one page limit and a tabular feature-oriented format for RFEX reports with components easily understood by our target users. These goals enable effective leveraging of powerful “human in the loop” concept for explainable AI as outlined in [4,26] where RFEX serves as “user interface” between ML and the users. Following the principles of *User Centered Design*, we first asked who our *persona* is, e.g. what are their goals, skills, pain points, frustrations, how do they expect/like to receive the results of explainability, and what questions they would like answered from ML explainers. Our typical persona is the domain expert, often the key decision maker for adoption of ML, with basic knowledge of statistical measures and familiar with typical formats of medical tests where a report contains a list of tests (analogous to features in ML), each usually with ranges or values expected for + or – cases. Their familiarity with the inner workings of ML systems is low to moderate only. Often they have their own set of test cases not used for ML training, which they use to establish trust (hence importance of ML Sample explainers as noted in [10]). They often have specific *explainability questions (EQ)* summarized below, where EQ 1-5 refer to *ML Model* and EQ 6-7 refer to *ML Sample explainability*:

1. What is the best accuracy achieved by the chosen ML method? (This is, in essence, the first and basic question addressed by most ML systems, black box or explainable)?
2. Which features (preferably small manageable number) are most important for predicting the class of interest (this questions is critical in reducing complexity of the problem space)?
3. What are basic statistics/ranges and separation of feature values between those for + and – class?
4. What is the tradeoff between using a subset of features and the related accuracy?
5. Which groups of features work well together?
6. Why was my sample classified incorrectly or correctly? Was it a “reliably” classified/misclassified sample or “marginal”?
7. Which features or factors contributed to incorrect classification of the sample?

### 2.1. RFEX Model Explainer

RFEX Model Explainer presented in this paper is a significant improvement of the original RFEX approach published in [16]. It provides one page *RFEX Model Summary* table using the following seven steps (steps 1-3 are the same as in [16], steps 4-7 are new). These steps directly answer users’ model explainability questions 1-5 as outlined above but use some novel, more standard and institutive measures than the original RFEX.

1. *<u>RF base accuracy</u>*: We train the RF classifier on a full training database using *all features* to establish baseline (ultimate) RF accuracy and optimal RF parameters e.g. *ntree, mtry and cutoff* using standard best practices for RF optimization. The accuracy measure we use is the F1 score.
2. *<u>Feature Rank</u>*: We rank features by their predictive power using Mean Decrease in Accuracy (MDA) measured from the trained RF classifier in Step 1. This step answers questions on which factors (features) are most important and how they rank in terms of their predictive power in RF classification. In the case of unbalanced training data (e.g. number of + class samples is 10% or less than – class samples), we recommend the use of class specific MDA (separate rankings for + and – class denoted by MDA+ and MDA-) which, in our experiments, clearly showed different ranking [16, 23] and is provided by the R toolkit [15].
3. *<u>Cumulative F1 score</u>*: We provide tradeoffs between using subsets of top ranked features (up to the topK) and RF accuracy by computing *“cumulative F1 score”* for each combination of top ranked 2, top ranked 3,…top ranked topK features (in each step we perform full optimization/training of the RF). We chose topK threshold such that the cumulative F1 score for topK features is close enough to base F1 score using all features (from Step 1). It is of significance that in multiple experiments we have done [16, 23, 24], a topK of only 2-6% of the total number of features produced over 90% of the accuracy achieved by using all features. This step drastically reduces complexity and dimensionality to about 10 topK features and allows RFEX summary reports to fit on one page, an important requirement for ease of use and simplicity.
4. *<u>AV/SD; [MAX,MIN] (class specific feature value ranges</u>):* To determine basic feature value statistics/ranges (e.g. presence or absence of a “signal” or a property, level of gene expression), we use common measures of average, standard deviation (AV/SD) and [MIN,MAX] range for feature values of + and - class, for each feature. This replaces feature Directionality (DIR) from [16] as a more direct, precise and familiar measure
5. *<u>Cohen Distance</u>*: To determine separation of feature values for + and – class for each feature, a common question from our target users, we converged on Cohen Distance (we prefer this vs. Mann-Whitney-Wilcox test due to the problem with p values for large data sets being very low and the property of Cohen Distance to indicate the degree of separation vs. only confirming the hypothesis [19]). Cohen Distance between feature values of two populations e.g. of + and – class is:

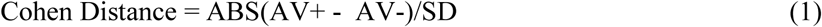

where *AV+* is the average of feature values for the positive class; *AV-* is the average of feature values for the negative class; and *SD* is the larger standard deviation of the two feature value populations. As noted in [19], Cohen Distance of less than 0.2 denotes a small; from 0.2 to 0.5 denotes a medium; from 0.5 to 0.8 a large, and above 1.3 a very large distance.
6. *<u>Cliques of N features</u>*: To determine feature interaction or most predictive groups of N features (clique of N) from the topK features (or sometimes from the 2*topK to increase the coverage), we perform full RF training for each combination (clique) and, for each, record their best F1 score. We show top (usually 10) cliques (e.g. list of features) with the best F1 scores. Our users were mainly interested in cliques of 3 and 4. Note that this exhaustive search is possible given the drastic feature reduction performed in Step 3. This measure replaces Mutual Feature Interaction (MFI) from [16] as a more accurate and intuitive measure computed directly using actual RF accuracy results. Note that top ranked N features may not necessarily form the best cliques of N.
7. *<u>RFEX Model Summary</u>*. We collect the data as a *one page tabular view* comprising of: a) a base RF accuracy data (from Step 1), e.g. F1 score, optimal ntree, mtry, cutoff; and b) an RFEX Model summary table with rows consisting of the topK features sorted by MDA (MDA+ for imbalanced training data case) and columns as in Steps 2-6 above, similar in format to common medical test reports. An example of RFEX Model Summary Table is shown in the case study in section 3.

RFEX Model Explainer summary is then used as explainability information on how RF works on the training data as a whole and can be used as a component of an “explanatory debugging” ML user interface as advocated in [26].

### 2.2. RFEX Sample Explainer

The RFEX sample explainability approach is designed with the target user in mind for situations when, for example, they want to check the RF classification results on a *sample or feature* level for a sample for which they know the ground truth, critical in forming users’ trust in ML systems [10]. Other use cases are in quality control (as in [22]) or editing of training data where one wants to delete samples of marginal “quality” in order to perform better ML training. RFEX Sample explainer consists of:

- *RFEX Sample Explainability Data* consisting of several *global sample level* types of information: a) *CORRECT_CLASS:* Correct (ground truth) class label of tested sample known or assumed by the domain expert; b) *RF_CLASS_LABEL*: Sample class label determined by RF; and c) *VOTE_FRACTION:* Fraction of RF trees (relative to *ntree* of trained classifier) voting for the CORRECT_CLASS, e.g. the ground truth class of the sample, using trained RF from Step 1 in section 2.1 in predict mode, with *all* the features. VOTE_FRACTION, when compared to the cutoff of the trained RF, helps user assess if the classification was done “reliably” or was “marginal” or wrong, and is provided in R [15].
- *RFEX Sample Summary Table* shows, in a one page summary, how *particular features* (topK features from RFEX Model Summary) contribute to RF decisions on tested samples. The nine columns of the RFEX Sample Summary table show all the information for easy analysis (the first five columns are the same as in RFEX Model Summary for easy reference) and are: *Feature Rank*; *Feature Name*; *Feature MDA rank*; *Feature AV/SD for positive class; [Min,Max]; Feature AV/SD for negative class; [Min,Max]; Feature Value of Tested Sample*; *Sample Cohen Distance to + class* measured as difference between sample feature value and average of feature values for + class, normalized by standard deviation [19]; *Sample Cohen Distance to – class*; and *K Nearest Neighbor Ratio* defined as:
  - *K Nearest Neighbor (KNN) ratio* **-** this measure looks at K nearest neighbors to current feature value and measures the fraction of those that belong to CORRECT_CLASS. It complements the *Sample Cohen Distances* in that it is more local and rank based, as well as non-parametric, and consistent with the ways RF forms decision tree boundaries. For K, we recommend 20% of the number of samples of the smaller class.

Examples of RFEX Sample Summary tables from our case study are shown in Tables 2 - 4. As with RFEX Model Summary, RFEX Sample summary is designed to facilitate “human in the loop” process by providing an easy-to-use format to help users in identifying problematic samples (e.g. outliers) *and* their problematic features. Domain experts then may follow with their domain knowledge for final decision making (e.g. keep the sample, reject it, drop some features). Users can interpret RFEX Sample summary by first analyzing its global data, referring to *sample as a whole* e.g. VOTE_FRACTION, to get an indication of how close or far from the vote cutoff the sample was (was it a “*reliable or high confidence*” classification or a “*marginal*” one). Other global measures or metrics of sample “quality” to be checked from RFEX Sample Summary table include: a) average of Sample Cohen Distances to the correct class (expected to be small e.g. 1 or less for reliably classified samples); b) average of Sample Cohen Distances to the incorrect class (expected to be larger, e.g. above 2); and c) average KNN, expected to be above 0.4 for reliably classified samples. To identify *specific features* that may have caused marginal sample quality (e.g. are “out of range”), one may apply measures as above to each feature, and possibly adjust the decision by feature importance (MDA rank). One can easily derive a set of rules using the above measures to interactively or automatically identify bad samples and/or bad features (similar to the tests on RF metrics developed in [22]).

**Table 1:**
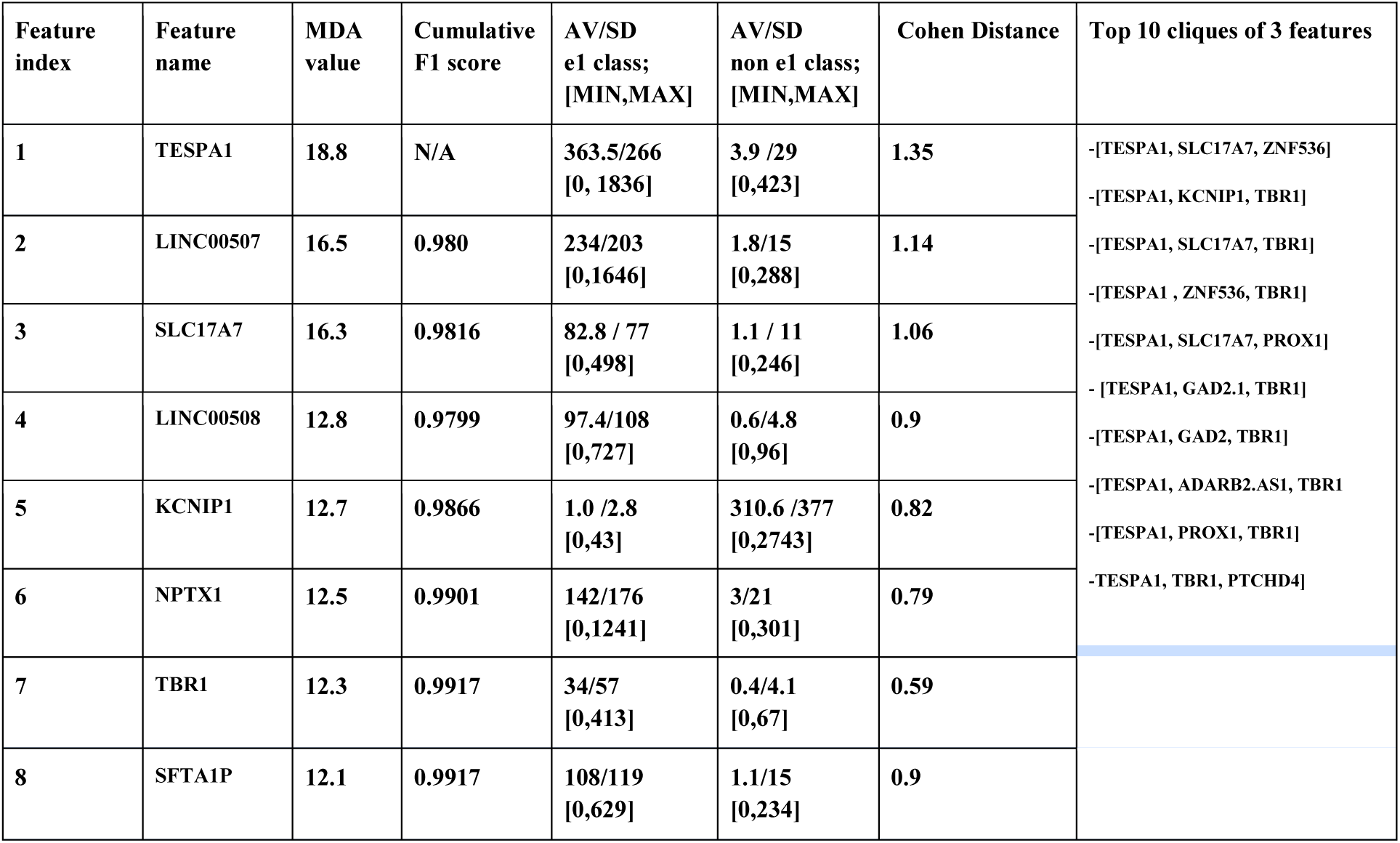
RFEX Model Summary table for JCVI data - e1 cluster. Base RF accuracy using all features is F1=0.995, for ntree = 1000, mtry = 50 and cutoff (0.3, 0.7)

**Table 2:** RFEX Sample Summary for JCVI data, Sample 1 – “good” sample, VOTE_FRACTION 100%

| Feature Rank | Feature name | Feature MDA rankings | Feature AV/SD for e1 (+) Class; Range [Min,Max] | Feature AV/SD for non-e1 (-) Class; Range [Min,Max] | Feature Value of tested sample | Sample Cohen Distance To e1 class | Sample Cohen Distance To Non e1 class | K Nearest Neighbor ratio |
| --- | --- | --- | --- | --- | --- | --- | --- | --- |
| 1 | TESPA1 | 18.8 | 363.5/266 [0, 1836] | 3.9 /29 [0,423] | 125.5 | 0.89 | 4.2 | 57/60 |
| 2 | LINC00507 | 16.5 | 234/203 [0,1646] | 1.8/15 [0,288] | 68.4 | 0.82 | 4.4 | 56/60 |
| 3 | SLC17A7 | 16.3 | 82.8 / 77 [0,498] | 1.1 / 11 [0,246] | 209.9 | 1.65 | 18.9 | 59/60 |
| 4 | LINC00508 | 12.8 | 97.4/108 [0,727] | 0.6/4.8 [0,96] | 14.1 | 0.77 | 2.8 | 49/60 |
| 5 | KCNIP1 | 12.7 | 1.0 /2.8 [0,43] | 310.6 /377 [0,2743] | 0 | 0.36 | 0.82 | 57/60 |
| 6 | NPTX1 | 12.5 | 142/176 [0,1241] | 3/21 [0,301] | 214.9 | 0.41 | 10.1 | 56/60 |
| 7 | TBR1 | 12.3 | 34/57 [0,413] | 0.4/4.1 [0,67] | 81.3 | 0.83 | 19.7 | 58/60 |
| 8 | SFTA1P | 12.1 | 108/119 [0,629] | 1.1/15 [0,234] | 126.5 | 0.16 | 8.36 | 56/60 |

**Table 3:** RFEX Sample Summary for JCVI data, Sample 2 – “marginal” sample, VOTE_FRACTION 80%. Possibly problematic features are marked ***

| Feature Rank | Feature name | Feature MDA rankings | Feature AV/SD for e1 (+) Class; Range [Min,Max] | Feature AV/SD for non-e1 (-) Class; Range [Min,Max] | Feature Value of tested sample | Sample Cohen Distance To e1 (+) class | Sample Cohen Distance To Non e1 (-) class | K Nearest Neighbor ratio |
| --- | --- | --- | --- | --- | --- | --- | --- | --- |
| 1 | TESPA1 | 18.8 | 363.5/266 [0, 1836] | 3.9 /29 [0,423] | 601 | 0.89 | 20.6 | 60/60 |
| 2 *** | LINC00507 | 16.5 | 234/203 [0,1646] | 1.8/15 [0,288] | 2 | 1.14 | 0.01 | 4/60 |
| 3 *** | SLC17A7 | 16.3 | 82.8 / 77 [0,498] | 1.1 / 11 [0,246] | 1 | 1.06 | 0.009 | 11/60 |
| 4 *** | LINC00508 | 12.8 | 97.4/108 [0,727] | 0.6/4.8 [0,96] | 1 | 0.89 | 0.08 | 31/60 |
| 5 | KCNIP1 | 12.7 | 1.0 /2.8 [0,43] | 310.6 /377 [0,2743] | 1 | 0 | 0.82 | 51/60 |
| 6 | NPTX1 | 12.5 | 142/176 [0,1241] | 3/21 [0,301] | 78 | 0.36 | 3.57 | 58/60 |
| 7*** | TBR1 | 12.3 | 34/57 [0,413] | 0.4/4.1 [0,67] | 0 | 0.6 | 0.1 | 31/60 |
| 8 *** | SFTA1P | 12.1 | 108/119 [0,629] | 1.1/15 [0,234] | 2 | 0.89 | 0.06 | 9/60 |

**Table 4.** RFEX Sample Summary from FEATURE data [16] for ASP_PROTEASE.4.ASP.OD1 for a marginal sample of + class, with VOTE_FRACTION of 70%. Possibly problematic features are marked with ***.

| Feature Rank | Feature Name | Feature MDA+ Rankings | Feature AV/SD for Positive class; [Min, Max] | Feature AV/SD for Negative class; [Min, Max] | Feature Value of Tested Sample | Sample Cohen Distance to Positive Class | Sample Cohen Distance to Negative Class | K Nearest Neighbor ratio |
| --- | --- | --- | --- | --- | --- | --- | --- | --- |
| 1*** | Solvent_Accessibility_s5 | 21.9611 | 291.9/112.3 [52, 1119] | 923.0/435.4 [7, 3547] | 1119 | 7.36 | 0.45 | 0/317 |
| 2*** | Secondary_Structure1_Is_Turn_s3 | 21.8895 | 8.3/2.0 [0, 14] | 1.78/2.7 [0, 22] | 3 | 2.63 | 2.34 | 24/317 |
| 3*** | Residue_Name_Is_Thr_s4 | 21.6433 | 5.7/1.7 [0, 12] | 0.85/1.6 [0, 14] | 3 | 1.59 | 1.38 | 108/317 |
| 4*** | Residue_Name_Is_Gly_s3 | 20.91 | 3.4/1.2 [1, 8] | 0.49/1.1 [0, 11] | 1 | 2.12 | 0.46 | 30/317 |
| 5*** | Solvent_Accessibility_s4 | 20.6998 | 167.8/94.2 [0, 593] | 673.4/361.9 [0, 3580] | 546 | 4.02 | 0.35 | 0/317 |
| 6*** | Secodary_Structure_1_Is_Strand_s5 | 20.6161 | 15.3/2.0 [5, 30] | 4.3/6.3 [0, 52] | 9 | 3.11 | 0.75 | 6/317 |
| 7*** | Residue_Class1_Is_Unknown_s3 | 20.5889 | 3.5/1.2 [1, 8] | 0.5/1.2 [0, 18] | 1 | 2.11 | 0.39 | 30/317 |
| 8*** | Residue_Name_Is_Gly_s2 | 20.4037 | 3.7/1.5 [0, 5] | 0.17/0.6 [0, 8] | 2 | 1.13 | 3.05 | 48/317 |

For RFEX Model and Sample explainer implementation, we use RF implemented in R because it is well supported and provides all the measures and features we need [15]. All RFEX steps can easily be automated in the form of a pipeline of processing steps, leveraging available RF toolkits like [15], as we demonstrated in the RFEX Model toolkit [20]. The RFEX output is in a standard and familiar tabular CSV format, which can be further customized as well as processed with “filtering” rules that are domain specific.

## 3. Application of RFEX to human nervous system cell type clusters from gene expressions using data from J. Craig Venter Institute and Allen Institute for Brain Science

In this case study we collaborated with the team of Dr. R. Scheuermann from J. Craig Venter Institute (JCVI), and used data (“JCVI data” in this paper) produced in a collaboration between JCVI and the Allen Institute for Brain Science as published in [21,27] to test and verify our RFEX methods. The goals of our case study were two-fold: a) investigate if RFEX Model Summary reflects information published in [21,27] in a correct and easy to use way, and b) use RFEX Sample explainer to identify *samples and features* that are possibly “out of range” and may need to be removed from the training set for quality control purposes (only based on the data not domain knowledge), an important use case in ML research and also done in [22]. JCVI data was an excellent test case for our evaluation since it contained some level of ground truth (e.g. identified most important gene markers for certain cell cluster types as in [21,27]), developed using domain-specific knowledge as well as RF methods. The matrix of gene expression values from single nuclei samples derived from human middle temporal gyrus (MTG) layer 1 forms the features for RF classification (610 of them), and groups them by 16 different cell type clusters, which constitute the classes for RF classification. The goal of researchers in [21,27] was to use RF analysis to identify small sets of gene markers (3-5) that define various cell type clusters and with this to contribute to the Cell Ontology (CL) database. They performed RF analysis to identify key gene sets that predict cell types, which served as ground truth information for our case study. We, as in [21,27], transformed the problem of 16 class classification into a set of 16 binary classifications, where all classes of non interest were grouped into the – class and the one cell type cluster (class of interest) was considered the + class. We analyzed several cell type clusters and in this paper we show RFEX analysis of cell type cluster *e1*. Cluster for cell type e1 had 299 positive samples and 572 negative (non-e1) samples, and 610 features (gene expression levels), and thus offered a relatively balanced data set. In [21], cluster e1 is defined as “A human middle temporal gyrus cortical layer 1 excitatory neuron that selectively expresses TESPA1, LINC00507 and SLC17A7 mRNAs, and lacks expression of KNCP1 mRNA”.

RFEX Model based explainer shown in Table 1 yielded results very consistent with the analysis in [21,27] and importantly pointed out some directions for further study. The whole analysis of RFEX explainability has been performed by using *only one page RFEX Model Summary* from Table 1, an important goal of the RFEX approach. To measure base RF accuracy with all 610 gene features (*explainability question EQ 1*), we performed a grid search using a range of RF parameters and obtained a very high F1 score of 0.995 with ntree = 1000, mtry = 50, cutoff of 0.7 for + class, 0.3 for – class (high accuracy RF classification for e1 cluster was also confirmed in [21, 22,27]). RFEX Model Summary Table 1 was then generated with only 8 topK ranked features achieving accuracy of F1 = 0.992, very close to the accuracy using all 610 features (F1=0.995), hence over 98% feature dimensionality reduction with practically no loss of accuracy. Cumulative F1 score also shows tradeoffs in using subset of features vs. achievable accuracy (EQ 2, 4). Importantly, the RFEX Model Summary table ranking (derived from 5 runs of RF and averaged) shows stable (low standard deviation) ranking [23] with the top 3 ranked genes *precisely matching* the e1 cluster definition from [21] (TESPA1, LINC005007, SLC17A7), with KNCNIP1, also mentioned in [21], ranked fifth (EQ 2). Stable MDA rankings were also by themselves one “confidence” indicator of whether RF was able to train well [23]. Furthermore, by observing AV/SD of class specific feature values in Table 1, one can easily find information about levels of gene expression e.g. top 3 ranked genes show high expressions (high AV for + class feature values, low AV for - class) and KCNIP1, quoted as predictive for its lack of expression in [21], shows low AV value for the + class vs. high AV for the – class (EQ 3). Cohen Distance values confirm that all features show good separation between + and – classes (average Cohen Distance of 0.94), and notably this separation is highest for highly ranked features (EQ 3). Cliques information measuring which groups of 3 features offer best prediction (3 was recommended for e1 in [21]), show 10 best cliques of 3 in the last column of Table 1 (EQ 5). Best cliques are dominated by TESPA1 and include highly ranked genes, but notably not all from the top 8 ranked ones. Interestingly and significantly, RFEX Model Summary Table 1 also pointed to some unexpected results that should be investigated, thus opening the possibility of gaining new knowledge from this kind of explainability analysis. First, there is number of highly ranked gene markers not mentioned in [21,27], with the most notable one, TBR1, also appearing in many cliques of 3 in Table 1. TBR1 raised interest for further study by our JCVI colleagues. As a quick check we used our gene interaction search and visualization tool GeneDive [25] and found that there are some published interactions between SLC17A7 and TBR1. We also achieved similarly consistent results for cell type cluster i1 [23].

In the second part of our case study on the JCVI data, we used our newly developed RFEX Sample Summary for a common case of quality control similar to [22]. We evaluated if it can help users identify samples that are “marginal” or “outliers”, and hence not recommended for use in ML training, as well as to identify which specific features among the topK highly ranked ones contributed to this. While in [22] this analysis is based partially on the use of domains specific information e.g. from the data extraction methods together with tests on derived RF metrics, our analysis is strictly based on RFEX Sample Summary data and not on any domain knowledge. From the JCVI e1 training data, we chose two possibly “edge” samples using VOTE_FRACTION as a first metric: a) *Sample 1 – reliably classified (good) e1 cluster sample* – with VOTE_FRACTION of 100% of trained RF trees using RF prediction engine on all 610 features treating this sample as a new sample; and b) *Sample 2 – possibly “marginal or outlier” e1 cluster sample*, with the least VOTE_FRACTION of 80% of trained RF trees. RFEX Sample Summary for Sample 1 is shown in Table 2, and for Sample 2 is shown in Table 3. Visual observation and simple analysis of Tables 2 and 3 (as recommended at the end of section 2) clearly points to differences between these samples (EQ 6). At the sample (global) level, the average of Cohen Distances to + class feature value population was 0.73 for Sample 1 and 0.74 for sample 2, but the average of Cohen Distances to - class samples was 8.66 for Sample 1 and only 3.16 for Sample 2. Importantly, average KNN ratio for Sample 1 was 0.93 (most neighbors of tested sample were of correct + class) but only 0.53 for Sample 2. These findings indicate much poorer “separation” of feature values for marginal Sample 2. Tables 2 and 3 also allow easy identification of specific features (genes) causing this marginalization using rules recommended at the end of section 2 (EQ 7). In Table 3 we marked with *** five such features (2,3,4,7,8) identified by: a) being closer to the wrong class (indicated by their Sample Cohen Distance values being smaller for the – class than to the correct + class), as well as b) having very low KNN (e.g. <0.4). To verify our hypothesis that the above five features (out of 610 features) are responsible for this sample receiving low votes and that lower VOTE_FRACTION indicates presence of “out of range” features” we replaced them with their respective average values for the + class and observed considerable increase of the VOTE_FRACTION from 80% to 93.7%. To further verify RFEX Sample Explainer and calibrate the “out of range” tests above, we also performed tests on Stanford FEATURE data [16] containing as features 480 electrochemical properties around molecular locations with target classes as functional sites (+) and non-functional sites (-). For functional model ASP_PROTEASE.4.ASP.OD1, the training data contained 1585 + (site) samples denoting active functional sites, and 48577 – (non-site) samples. RF was able to achieve F1 score of 0.999 [16]. RFEX Sample Summary in Table 4 shows data for a marginal + class sample (one receiving the smallest number of trained RF tree votes among all + samples), which had VOTE_FRACTION of 70%. Data for this sample show a very similar “pattern” as for JCVI data, with the average of Sample Cohen Distances to + class of 3.0, average of Sample Cohen Distances to the - class of 1.15 (e.g. sample features are closer to the wrong – class than the + class) and average KNN of 0.097, all well below expected values for “good” samples. As a reference, for the + sample in this database, with VOTE_FRACTION of 100%, the average of Sample Cohen Distances to + class was 0.36, average of Sample Cohen Distances to the - class was 2.68, and average KNN was 0.47. Similar tests as for JCVI data on individual features of FEATURE marginal sample identified all eight of them being “out of range”. To measure relation of VOTE_FRACTION to the presence of “out of range features” we “corrupted” good sample’s top 4 feature values by replacing them with respective averages of feature values for – (wrong) class and got significant drop in VOTE_FRACTION from 1.0 to 0.74.

## 4. Discussion

In this paper we presented novel RFEX Model and Sample explainers for RF classifier. RFEX is designed from the ground up with non-ML experts in mind, and with simple and familiar formats and components e.g. providing one-page tabular outputs and measures familiar to most users. In this paper we presented: a) significant improvements in *RFEX Model explainer* compared to the previously-published version; b) a new *RFEX Sample explainer* that provides explanations of how RF classifies a particular data sample and designed to directly relate to the RFEX Model explainer; and c) an RFEX Model and Sample explainer case study on the data and findings from J. Craig Venter Institute (JCVI) and Allen Institute for Brain Science. In our case study we demonstrated that RFEX showed correct and useful results in a format that is simple and familiar to our target users, as well as helped to explain RF classification at overall model as well as at sample and feature level, important functions for increasing users’ trust and for quality control in ML training. We also showed that RFEX could even identify some new areas (e.g. genes) for further investigation. RFEX is easy to implement using available RF toolkits as we demonstrated, and its one page tabular format offers easy to understand representation for non-experts. Future work includes more testing of RFEX with users and on data where sufficient ground truth (including at sample and feature level) is available, and completing full function RFEX Model and Sample Jupyter based toolkit.

## 5. Acknowledgements

We are grateful to a number of people who made this work possible: Dr. R. Scheuermann and B. Aevermann from JCVI for the data for our case study, their feedback and guidance; Prof. Russ Altman for years of encouragement and support, and for general feedback on RFEX and FEATURE data; Mike Wong from SFSU CCLS for feedback on RFEX as it applied to FEATURE data; and Dr. Les Kobzik for constructive suggestions. Work has been partially supported by NIH grant R01 LM005652 and by SFSU COSE Computing for Life Sciences.

